# Identification and characterization of a biphasic/bidirectional wound-inducible *RHA3B* gene promoter from *Arabidopsis thaliana*

**DOI:** 10.1101/2021.01.28.428589

**Authors:** Venkateswara Rao, Vijaybhaskar Virupapuram

**Affiliations:** CSIR-Centre for Cellular and Molecular Biology (CCMB) Habsiguda, Hyderabad-500007

**Keywords:** *Arabidopsis thaliana*, DsE enhancer, Wound induction, Promoter, RHA3B, MRPL11

## Abstract

Conditional promoters such as wound inducible are important tools for plant biotechnology to selectively express agronomically important genes. From a set of Enhancer trap (DsE-uidA) transposant lines we identified wound inducible ET1075 line by injuring leaf blade and screening for Ds linked GUS gene expression. Earliest GUS stain detected at petiolar region at 2hrs post injury with a progressive attainment of maximum visual intensity between 12 and 24hrs wherein, the transcript expression induced in a bidirectional manner. DsE was found to be inserted in the intergenic region between *AT4G35480* and *AT4G35490*. To find the essential transcriptional regulatory region, deletion constructs comprising upstream sequences of p*RHA3B* fused to GUS reporter gene were functionally tested in Arabidopsis plants by generating stable transgenics. A 481 bp of upstream sequence from ATG is found to be sufficient to promote wound inducible gene expression. Sequence analysis revealed the presence of several regulatory elements implicated in wound inducible gene expression. Comparative analysis with similar wound inducible promoters revealed the presence of common *cis*-regulatory elements. This promoter can essentially be used in pest control and in molecular pharming to conditionally express agronomically/commercially important genes in plants.

**Highlights:**

- DsE enhancer trap transposant lines were generated.
- Identified a novel wound inducible promoter line ET1075.
- Wound inducible promoter p*RHA3B* is bidirectional and induces the expression of *RHA3B* and *MRPL11* genes.
- Wound responsive key cis-elements WRKY, W-box, FORC^A^ are present in the p*RHA3B* promoter.

## 1. Introduction

Promoters are the key regulators of gene expression at the transcriptional level. The choice of specific promoters used within a transgene construct is crucial to achieve the desired transgene expression in the temporal, spatial and measurable manner. Constitutive, tissue specific and conditional promoters are powerful molecular genetic tools to express foreign genes in plants. Constitutive promoter CaMV35S has been widely used in plants to express genes ubiquitously [1]. Such other promoters identified respectively in monocot and dicots are pActin and pMtHP [2,3]. The effect of ubiquitous gene expression is often harmful to the health of the plant by negatively effecting the desired output although such promoters are desirable to express several genes. One such example is DREB1A when over expressed under the control of CaMV35S promoter the transformants suffered growth abnormalities [4]. Hence, it is desirable to use tissue specific promoters over constitutive gene expression whose site of action/expression is of paramount interest. Some examples of reported tissue specific promoters are; Root specific p*GLT* [5], Leaf specific p*RbcS* [6], stem specific promoters p*DIR16*, p*OMT* [7], Flower specific p*CHS* [8] and Seed specific Napin promoter [9]. Chemically regulated inducible promoters reported are Cu repressible p*CYC6* promoter [10], the ethanol inducible *AlcA* promoter [11]. Within this group, there are promoters modulated by abiotic factors such as light, heat and cold.

Conditional or inducible promoters are another class of promoters that offer additional advantage of controlling gene expression both in tissue specific and in temporal manner. Wound inducible promoters can ideally be induced quickly and easily, preferably without the application of potentially expensive or toxic chemical inducing agents. Several wound inducible genes have been characterized by functional analysis of their promoter region in transgenic plants, including of those encoding proteinase inhibitors [12], phenylalanine ammonia-lyase [13], pathogenesis-related proteins [14], nopaline synthase [15]. The most desired strategy for pest control would be to express these genes of interest from a promoter that would express only at the site of feeding by larvae. Studies have identified few wound inducible promoters such as mannopine synthase 2’ (*mas2’*) promoter element from the T-DNA of *Agrobacterium tumefaciens* that was found to direct wound-inducible and root-preferential expression of a linked *uidA* gene in transgenic tobacco plants [16], wound-inducible promoter wun1 from tobacco [17] and a pathogen/wound-inducible PR10 promoter from *Pinus monticola* [18].

Although several tissue specific promoters are identified, very few wound inducible promoters are reported till date. In this study we report the identification and characterization of a *RHA3B* (RING-H2 FINGER A3B) gene promoter from *Arabidopsis* of which 482 bp is sufficient to confers wound inducible expression. The promoter sequence identified in this study has potential for application in selective expression of genes of interest.

## 2. Materials and methods

### 2.1 Plant material and growth conditions

*Arabidopsis* ecotypes Col-0 plants were used in the present study. The plants were grown at 21°C with light intensity of 8000 Lux and a 16 h/8 h light/dark cycle.

### 2.2 Identification of wound inducible Enhancer trap line

We employed enhancer detection screen to identify genes that are expressed specifically upon injury to leaf blade using serrated edged stainless steel forceps. A collection of approximately 180 independent enhancer trap lines of *Arabidopsis* were generated according to the protocol described [19]. The enhancer trap construct carried an engineered Ds transposon bearing a minimal promoter driven *uidA* (GUS) reporter gene whose expression is not detectable without being inserted in the vicinity of an enhancer. The enhancer trap lines at rosette stage of the plant were manually wounded by gripping leaf blade for 6 seconds with a forceps and were stained for GUS gene expression after 8 hours post injury.

### 2.3 Southern blotting and identification of DsE insertion in ET1075 line

For Southern analysis of the ET1075 line, plant genomic DNA was isolated according to the Cetyl-Trimethyl Ammonium Bromide (CTAB) method. Approximately 4 *μ*g of genomic DNA was digested with *Eco*R1 enzyme, electrophoresed and blotted onto a N^+^ nylon membrane and probed using randomly radio-labeled transposon sequences and auto-radiographed as described [19]. Thermal asymmetric interlaced (TAIL)-PCR was carried out to identify the DsE insertion sequence using the degenerate primer AD2 in combination with the nested primers Ds5-1, Ds5-2 and Ds5-3 for primary, secondary, and tertiary amplification reactions, respectively [20]. Genomic DNA from the promoter region of *At4g35480* was amplified using the primers designed to contain appropriate restriction sites (*Hind* III and *Sal*1), sub-cloned into the pMOS-Blue (Amersham Pharmacia) blunt end cloning vector following the manufacturer’s instructions and the insert DNA was sequenced using Ds5-3 and M13F primers.

### 2.4 Full length Promoter and deletion constructs

In order to find if the Ds tagged intergenic region between *AT4G35480* and *AT4G35490* is wound inducible, entire 3.12kb upstream sequence of RHA3B (*AT4G35480*) gene along with a part of its coding sequence was amplified using 3120FP and ComnRevP primers and cloned in promoterless vector pBI101.2 to make pRHA3B-FL::GUS fusion reporter construct. Further to find minimal wound inducible upstream sequence of *RHA3B*, three 5’ deletions pRHA3B-S, pRHA3B-M and pRHA3B-L were made using Hind III engineered forward primers (0515FP, 1057FP and 2086FP) with a common reverse primer (Comn RevP containing Sal1 site). PCR amplified DNA fragments were cloned in to promoterless pBI101.2 binary vector to obtain promoter GUS fusion constructs. Proper orientation was confirmed using GUS upstream reverse primer (GUS Ups) and 0515FP primers (Supplementary table 1).

### 2.5 Plant transformation and analysis of GUS expression

Promoter: GUS constructs in the binary vector pBI101.2 were mobilized into the *Agrobacterium* strain AGL1 by triparental mating using *Escherichia coli* HB101 (pRK2013) as helper. Transformation of Arabidopsis *in-planta* was carried out using the method described [21]. Transformants were selected by planting seeds on MS plates containing 2% Sucrose and 50 mg/l Kanamycin. GUS expression was analyzed after wounding as described [19].

### 2.6 cDNA synthesis and Real Time PCR analysis

To investigate the transcription levels of *RHA3B*, mitochondrial ribosomal gene L11 (*MRPL11*) flanking the DsE insertion, real time PCR analysis was done. For RT-PCR, independent WT Columbia plant leaf samples (unwounded/wounded) were collected from the rosette at 0, 0.5, 1, 2, 4, 6, 8, 12, 16 and 24 hours post injury. Total RNA was isolated from each plant sample using Trizol (Invitrogen) according to manufacturer’s protocol. Equal amounts of RNA (1μg) from different samples were reverse transcribed using SuperScript II RT (Invitrogen) according to the manufacturer’s instructions. RT-PCR was done by using the 7900 HT Fast Real time PCR system [ABI]. *GAPC* was used as endogenous control. With housekeeping gene GAPC, the relative amount of the *RHA3B*, ribosomal gene L11 transcripts in the different samples was calculated according to the comparative threshold 2^−**ΔΔCT**^ method [22].

## 3. Results and Discussion

### 3.1 Identification of wound inducible promoter through DsE enhancer trap lines

To identify the wound inducible promoter, 180 independent enhancer trap lines were generated according to the described method [19]. Out of the 180 lines screened one-line ET1075 was found to show wound-inducible gene expression, 8 hours post injury to leaf blade (Fig. 1A). To assess the copy number of the transposon in ET1075, F3 seeds were germinated on plates containing kanamycin. Out of 56 seedlings tested, 40 were kanamycin resistant (KanR) and 16 were sensitive to kanamycin (KanS). The KanR:KanS ratio was not significantly different from a value of 3:1, which would be expected for a single-copy insertion (0.18>*P*>0.1). Southern analysis of *Eco*R1 digested genomic DNA of the line probed with randomly radio-labeled DsE construct revealed as expected 3 bands i.e., one internal 2.7 kb and two junction fragments of 10 kb and 3 kb in the developed autoradiogram, confirming the single copy DsE insertion in the genome (Fig. 1B). To determine the site of insertion in the genome, TAIL-PCR was done on genomic DNA of ET1075 using Ds5-1, Ds5-2, Ds5-3 and AD-2 arbitrary degenerate primer to amplify the 5’-flanking sequence of the insertion. A 70 bp down ward shift of ~1 kb tertiary band was seen to that of ~1.1 kb secondary TAIL-PCR product suggesting that the amplicon originated from the same locus (Fig. 1C). Amplified product from the Secondary TAIL-PCR was cloned in to pMOS-Blue vector and the DNA was sequenced using M13F and Ds5-3 primers independently. Sequence analysis of the secondary TAIL-PCR product indicated identity to a portion in the intergenic region between *RHA3B* (*AT4G35480*) and *MRPL11* (*AT4G35490*) genes on chromosome number 4 (Fig. 1D, Supplementary Fig. S1). Two genes are separated by a 3082 bp and are transcribed in opposite directions. Exact insertion is at 2066 bases upstream to *AT4G35480*. *AT4G35480* encodes a putative RING-H2 finger protein RHA3B and *AT4G35490* encodes a mitochondrial ribosomal protein L11 (MRPL11). As some ring finger genes are involved in gene regulatory function, we decided to characterize *RHA3B* gene promoter if it conferred wound inducible gene expression.

**Figure 1.**
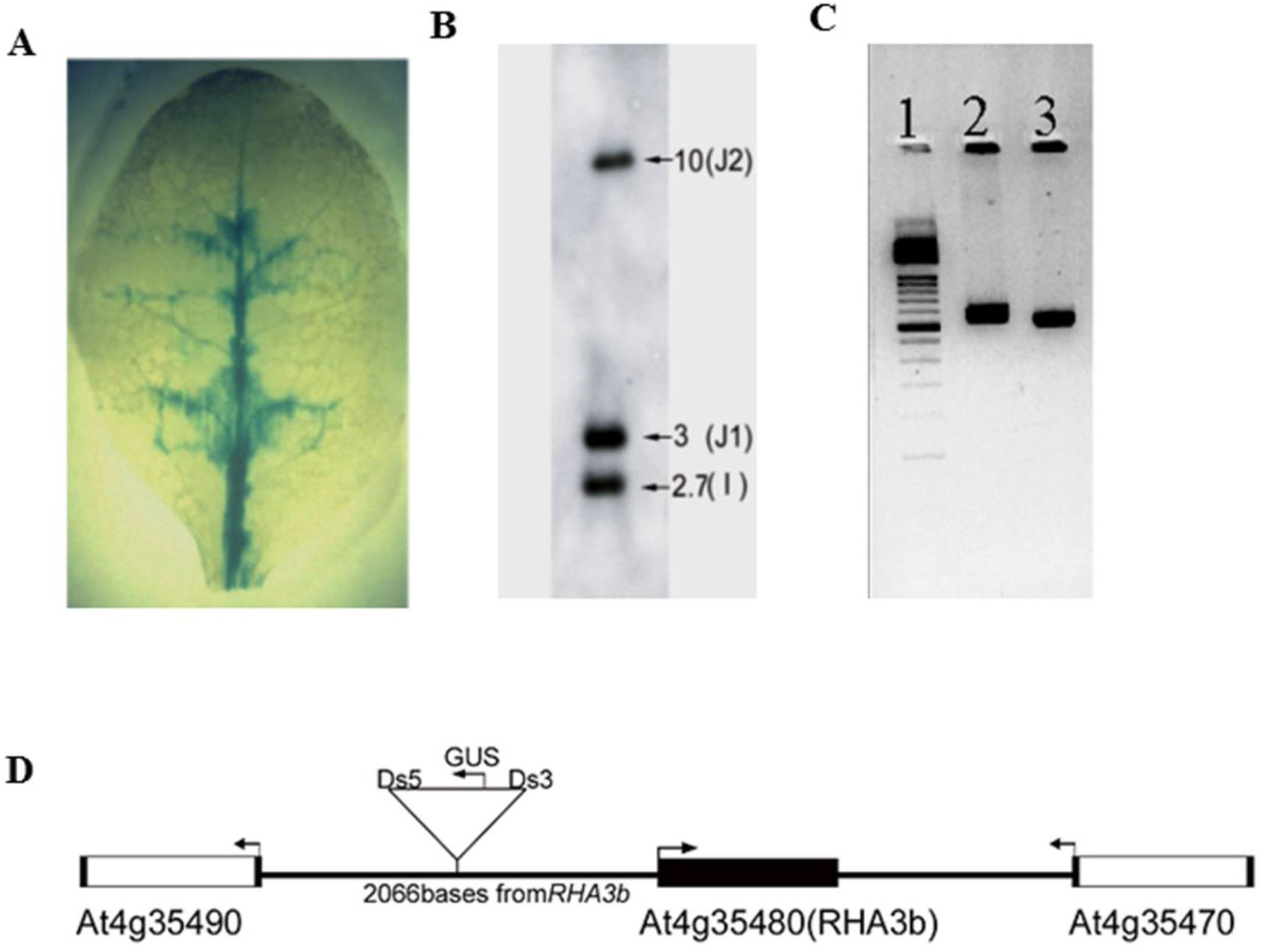
Characterization of the Arabidopsis Enhancer trap line ET1075. A) GUS stained rosette leaf of Enhancer trap line, ET1075. B) Southern blot of *Eco*R1 digested genomic DNA of ET1075 probed with DsE transposon. 2.7 kb band represents internal fragment (I) and 3 and 10 kb bands represent two junction fragments (J1 and J2) indicative of a single copy insertion. C) TAIL-PCR on ET1075 genomic DNA. Lanes 1: Marker, Lanes 2, 3: secondary and tertiary TAIL-PCR products, respectively D) Schematic diagram showing the insertion of DsE in between *RHA3B* and *MRPL11*.

### 3.2 Wound induced expression analysis of enhanced trap line ET1075

To study wound inducible expression in detail, ET1075 plant leaves at rosette stage were wounded and GUS stained at 2h, 4h, 6h, 12h, 24h, 26h, 36h, and 60 hrs post injury. GUS staining/gene expression was detected as early as 2h after injury and intensity of GUS stain gradually increased predominantly in the midrib and in branched venation. Maximum expression is attained between 12h and 24 hrs post injury (Fig. 2B). Site of insertion in the ET1075 was determined to be in the promoter region of *RHA3B* gene (Fig. 1D). To observe specific gene *RHA3B* expression at transcript level, reverse transcription was done on total RNA isolated from the wounded and unwounded leaves (12hrs post injury). *GAPC* was included in the RT reaction as the positive control. A downward shift of *GAPC* amplicon banding pattern would be seen in comparison to the genomic DNA amplicon for the same primers if cDNA is properly made. RT PCR analysis indicated that there was a basal level of expression of *RHA3B* in unwounded leaves, which was further triggered to higher level of transcription upon injury to leaf blade (Fig. 2A).

**Figure 2.**
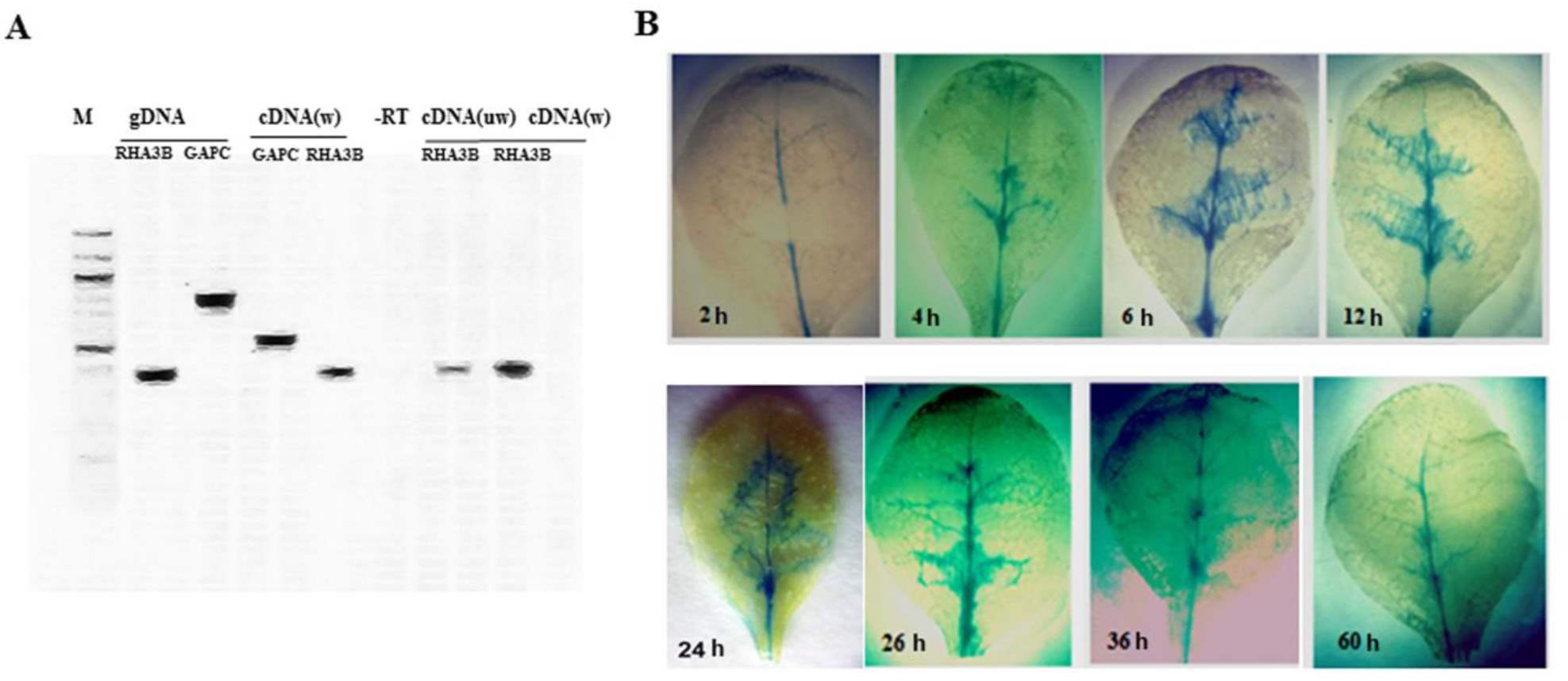
Wound induced gene expression analysis of ET1075. A) RT PCR analysis of DsE tagged RHA3B gene. Lane 1; Marker(100bp), Lanes2,3; genomic DNA amplified by RHA3B and GAPC gene specific primers, respectively. Lanes 4,5; cDNA of wounded plant amplified with GAPC and RHA3B gene primers, respectively. Lane6; No template control Lanes7,8; cDNA of unwounded and wounded plants amplified with RHA3B specific primers. B) GUS expression pattern of the tagged promoter upon wounding at various time intervals (h=hours).

### 3.3 Identification of the promoter region essential for wound inducible expression

Initially to verify if the site of DsE insertion contained wound inducible regulatory region, entire intergenic region (IGR) of 3.12 kb upstream to *RHA3B* along with 37 bases of coding region was amplified and cloned in-frame with the *uidA* (GUS) gene in binary vector pBI101.2 and transformed in to Arabidopsis plants. Transformed plants selected on Kanamycin were screened for wound induced expression upon injury. 60% (15/25) of the plants displayed wound inducible GUS expression pattern suggesting that *RHA3B* promoter confers wound responsive gene expression (Fig. 3B). To further define the promoter region required to confer wound inducibility three promoter deletion constructs pRHA3B-S, pRHA3B-M and pRHA3B-L were made comprising 481 bp, 1027 bp and 2057 bp respectively along with 37 bp exonic region to make in-frame translational fusion with GUS gene (Fig. 3A). Three deletions were amplified using previously cloned 3.12 kb target DNA and cloned to pBI101.2 binary vector and transformed to Arabidopsis Col-0 plants. 30 Kan^R^ transgenics for each of pRHA3B-S and pRHA3B-M and 23 for pRHA3B-L were obtained. When screened for Gus expression upon injury, 73% (22/30) of pRHA3B-S, 66% (20/30) of pRHA3B-M and 65% (15/23) of pRHA3B-L transgenics were found to be wound inducible (Table 2). As 73% (22/30) of pRHA3B-S plants exhibited wound induced GUS expression suggests that the promoter DNA sequence as small as 481 bp is sufficient to trigger wound inducible gene expression (Fig. 3B).

**Figure 3.**
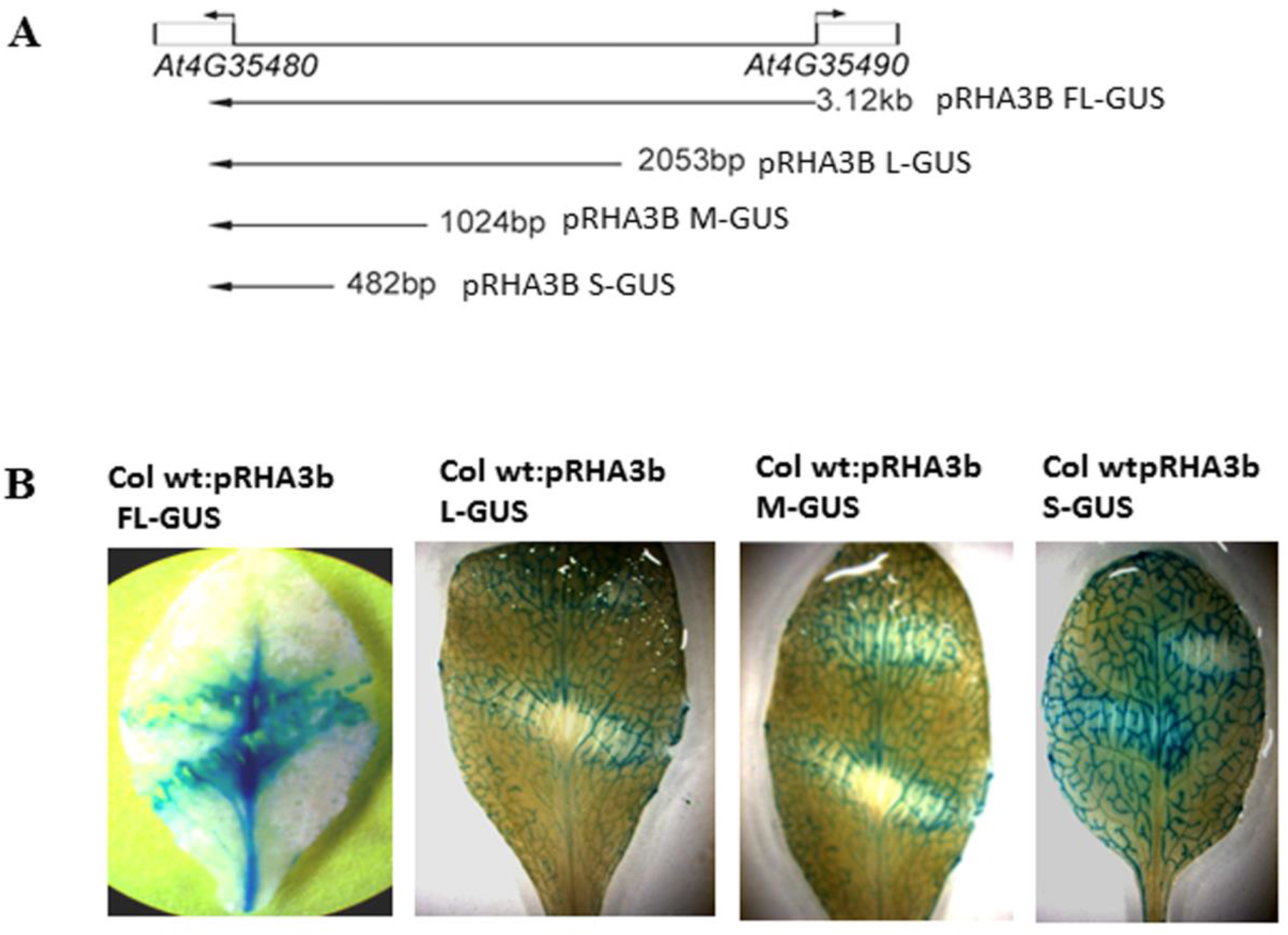
Identification of the promoter region essential for the wound induced expression. A) Schematic diagram showing the deletion constructs made to define the essential regulatory region. Full length pRHA3B FL-GUS comprising 3.12 kb entire intergenic region between AT4G35480 and AT4G35490 genes and three 5’-promoter deletion constructs pRHA3B-L, pRHA3B-M and pRHA3B-S comprising 2057 bp, 1027 bp, and 481 bp, respectively are depicted. B) Images showing wound induced GUS expression in transgenic plant leaves of full length and 5’deletion constructs pRHA3B:L-GUS, pRHA3B:M-GUS and pRHA3B:S-GUS.

### 3.4 Wound induced expression analysis of RHA3B and MRPL11 genes

To study the *RHA3B* gene expression in detail triggered by wound, real time PCR analysis was done. The fold change in expression of the target gene (*RHA3B*) was studied relative to the endogenous control gene (*GAPC*). Experimental results suggested that the *RHA3B* transcript expression induced to a level of 72, 103, 7, 1.45, 7.74, 2.62, 9.87, 0 and 0 fold corresponding to 30min, 1h, 2h, 4h, 6h, 8h, 12h, 16h and 24 hrs post wounding respectively (Fig. 4A). Gene transcripts were induced by wounding within 30 min and peaked at 1 h after wounding indicating that *RHA3B* is an early wound inducible gene. Enhancer trap *DsE* element inserted in intergenic region between *AT4G35480* and *At4G35490* could be influenced by the relevant *Cis*-elements on either side of the insertion. Hence, we looked for the status of adjacent gene *AT4G35490* (*MRPL11*) expression by real time PCR if it was influenced by the wounding. *MRPL11* expression induced to a level of 2, 1.55, 1.85, 5.45, 2.44, 5.5, 1.83, 7.18 and 12.67 fold at 30 min, 1h, 2h, 4h, 6h, 8h, 12h, 16h and 24 hrs post wounding, respectively (Fig. 4B). On contrary to *AT4G35480* whose gene expression was higher immediately after wounding and not detected at 16h and 24 hrs post wounding, *AT4G35490* transcript induction was slow and attains 7 and 12 fold at 16h and 24hrs post wounding. This data suggests both the gene *RHA3B* and *MRPL11* transcripts induced to several fold upon wounding and the pattern of their expression is bidirectional.

**Figure 4:**
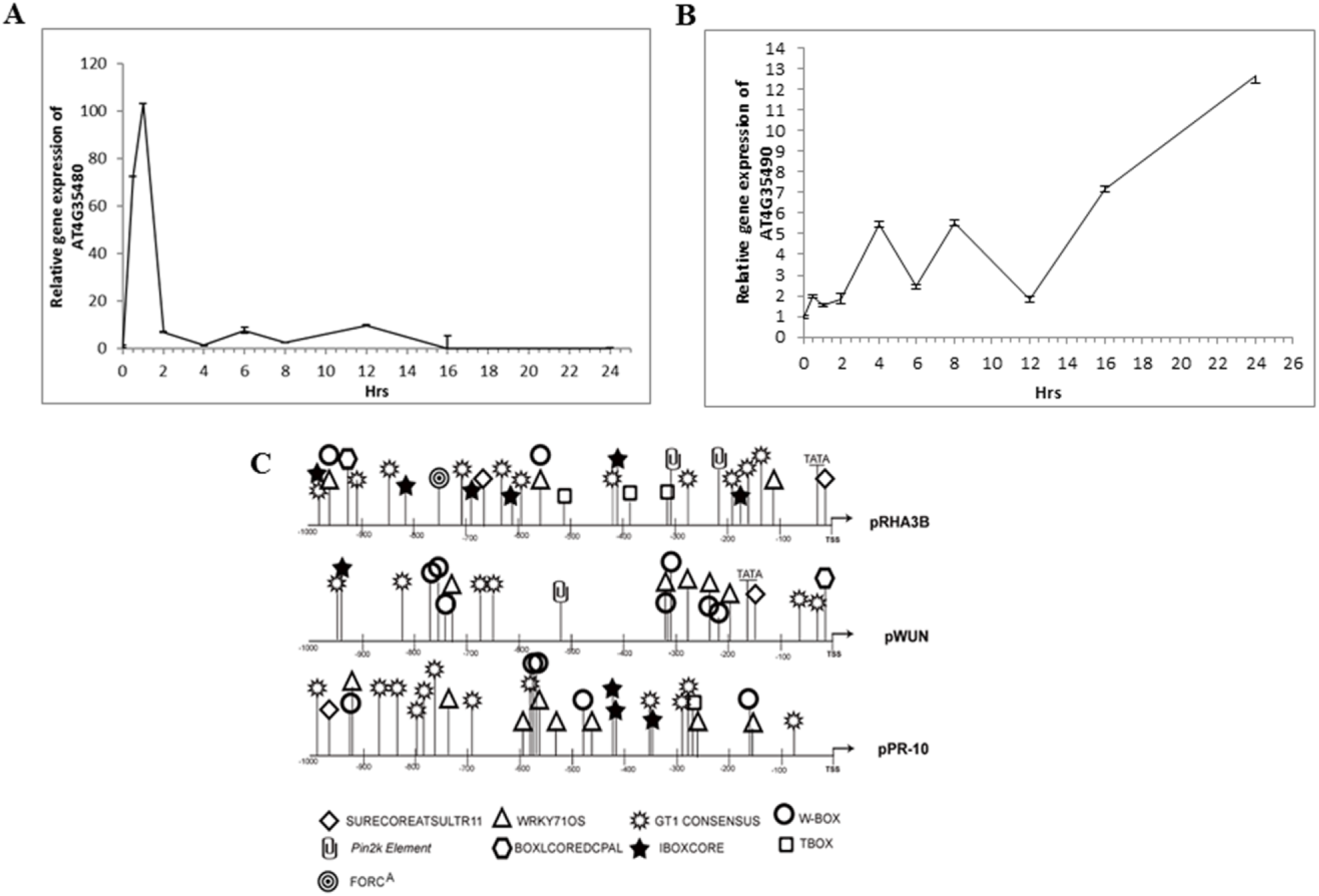
Wound induced expression analysis of *RHA3B* and *MRPL11* genes and comparison of Cis elements in the wound induced promoters. Relative quantities of A) *RHA3B* and B) *MRPL11* mRNA at various time points post-injury. The *GAPC* gene was used as internal control. Values shown represent the mean reading from three biological replicates and the error bars indicate the standard error of the means. C) Comparison of Cis acting elements of wound inducible promoters pRHA3B, pPR10 and pWUN. Several essential cis elements associated with wound induction are found across the pRHA3B, pPR10 and pWUN promoters such as WORKY71OS, W-Box.

*Cis*-acting elements present in the promoter region of a gene are required for gene expression/regulation. In order to see if the Cis acting elements are conserved across wound inducible promoters, 1kb upstream DNA sequence from the transcription start site (TSS) of pRHA3B along with two other wound inducible promoters pWUN, pPR10 [17,18] were analyzed using Plant Cis-acting Regulatory DNA Elements (PLACE) database [23]. Apart from TATA and CAAT common *Cis*-elements found in 1 kb pRHA3B; one site each for BOXLCOREDCPAL and FORCA elements, 2 sites each for BIHD1Os, MYCCONSENSUSAT, Pin2K, SURECORESULTR11, TBOX and WBOX elements, three sites for WRKY71OS, 8 for IBOX, and 15 for GT1CONSENSUS were found. All these cis elements are conserved among the three wound inducible promoters (Fig. 4C). BOXLCOREDCPAL [ACCWWCC] is required for the activation the Carrot Phenylalanine Ammonia-lyase gene (DcPAL1) in response to elicitor treatment [24]. *FORC*^A^ (T/ATGGGC) is a conserved hexameric promoter motif found in Arabidopsis genes co-expressed in response to fungal pathogens as well as defined light treatments [25]. BIHD1Os [TGTCA] induction is associated with resistance response in rice [26]. Pin2K element (AGTAAA) deletion is associated with loss of wound inducible gene expression [27]. A motif AGATCCCA similar to AG-motif AGATCCAA, sufficient to confer responsiveness to wounding and elicitor treatment is present at −1314 from TSS [28]. WRKY TFs are known to be wound response regulatory factors. W box [(T)TGAC(C/T)] is implicated in elicitor triggered response. So, the presence of several cis-acting elements contributes to the complex yet unique expression profile of a particular gene. In conclusion, we have screened a set of DsE Enhancer trap lines containing GUS reporter and found a novel wound inducible promoter of a Ring finger *RHA3B* gene. Identified promoter is bidirectional and induces the gene expression of *RHA3B* and *MRPL11*. Deletion analysis revealed 481 bp upstream sequence from ATG sufficient to confer wound induced gene expression upon mechanical injury. Discussed known Cis-regulatory elements might be contributing to the wound inducible expression of *RHA3B*. The isolated promoter could be used to engineer agronomically important conditional gene expression in crop improvement.

## Acknowledgements

We express gratitude to Dr. Imran Siddiqi for providing us the Gene trap and Enhancer trap lines. We gratefully acknowledge the funding from CSIR-India.

## Supplementary data

**Supplementary Figure S1.**
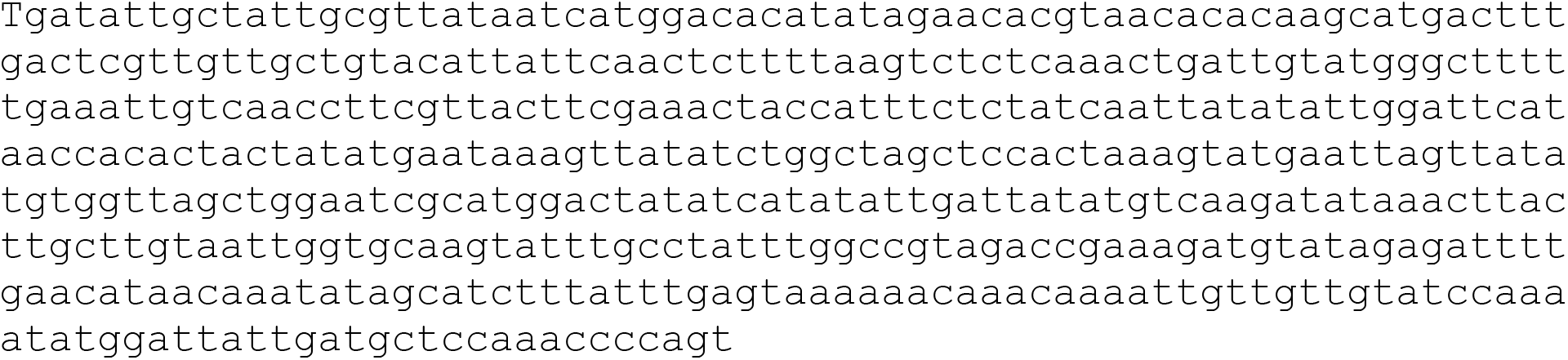
TAIL-PCR Clone sequence of DsE enhancer trap line ET1075. The sequence is located at Chr.4 16855379— 16854901 (Intergenic region between At4g35480 and At4g35490 of Chromosome no.4)

**Supplementary Table 1.**
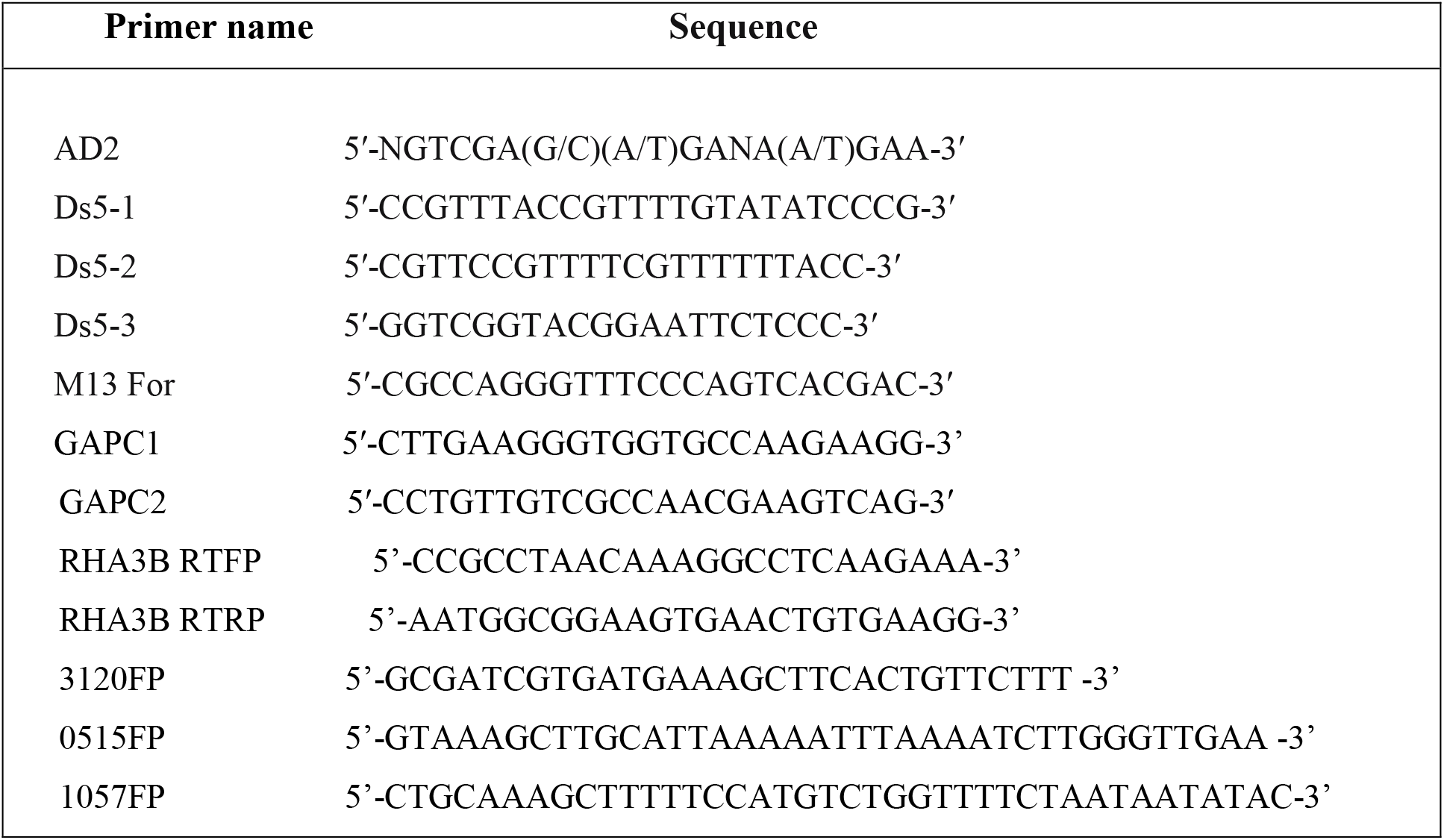
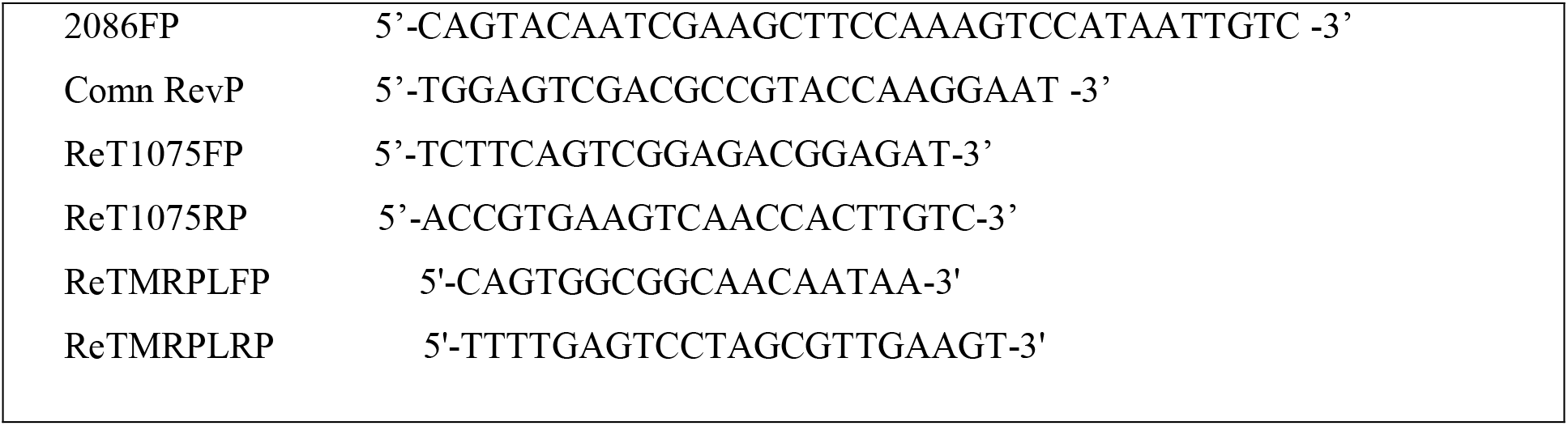
Primers used in the present study.

**Supplementary Table 2.** Wound induced GUS expression pattern in Transgenic Arabidopsis plants.

| **GUS expression in T1 transgenic plants** |  |
| --- | --- |
| **Promoter**<br>**(size in bp)** | **No. of Positive lines**<br>**for GUS staining** |
| pRHA3b-S (515) | 22/30 (73%) |
| pRHA3b-M (1057) | 20/30 (66%) |
| pRHA3b-L (2086) | 15/23 (65%) |
| pRHA3b-FL (3120) | 15/25 (60%) |
| Vector control | 0/30 |

